# Neural correlates of reward anticipation in 2000 children aged 9-10 years: relation to psychotic-like experiences and depressive symptoms

**DOI:** 10.1101/421826

**Authors:** Lisa Ronan, Graham K. Murray

## Abstract

**Background:** Schizophrenia spectrum disorders and depression have been associated with reductions in brain activation during reward anticipation. It is not known whether brain signals associated with reward anticipation relate to psychopathology dimensions of depression or schizophrenia in childhood prior to adolescence.

**Method:** We examined whether fMRI brain correlates of reward anticipation related to psychotic-like experiences and symptoms of depression, in 2129 children from the ABCD study aged 9-10 years.Psychotic-like experiences and depression were assessed using the Prodromal Questionnaire Brief Child version and the K-SADS. We fused regional MRI summary statistics for reward anticipation activation in the ABCD study data release 1.0 (contrast of expected large reward versus neutral expectation). Relations between brain activation and psychopathology were assessed using linear regressions in R for 82 brain regions, corrected for multiple comparisons for the number of regions using false discovery rate.

**Results:** From several regressions, there was an isolated unilateral association between right parsorbitalis activation and psychotic-like experiences, but no other significant associations between brain activation and psychopathology.

**Conclusions:** In 9-10 year old children, reward anticipation is not strongly related to psychotic-like experiences or depression. As previous evidence links depression and schizophrenia to reduced reward anticipation in adults and older adolescents, it appears likely that such associations develop over the adolescent period: this can be tested in follow-up studies of the ABCD cohort.

## Introduction

Dysfunction of the neural circuits underpinning reward processing has been hypothesised to relate to the pathogenesis of the symptoms of schizophrenia (1)(2)(3). Studies in adult patients with schizophrenia have indicated reduced brain activation during reward anticipation, especially in the ventral striatum (4)(5)(6)(7). The Monetary Incentive Delay (MID) task (8) is the functional MRI paradigm that has been used most extensively in both health and mental disorder to probe the neural basis of reward anticipation. Within patients with schizophrenia, inter-individual variability in the degree of reduced striatal activation on the MID has been associated with the severity of negative and depressive symptoms (6)(7), consistent with the possibility that ventral striatal dysfunction in schizophrenia compromises reward anticipation, leading to real world deficits in motivation and/or enjoyment in this disorder. A recent study, however, found no evidence of ventral striatal dysfunction in help-seeking young adults with prodromal symptoms of psychosis (9). However, reduced brain reward anticipation in the MID has also been shown in in other disorders, and may relate to severity of depressive symptoms in a transdiagnostic fashion (5)(6).

Thus far, the majority of clinical studies of reward anticipation using fMRI have examined older adolescents and adults. However, the largest study to date of the MID has been the IMAGEN study of around 2000 14 year olds (10). This study showed the MID task robustly activated a large network of brain regions, including the ventral stratum and medial prefrontal cortex, in addition to several other regions, as has been shown previously in adults. In IMAGEN, current clinical depression (n=22) or sub threshold depression (n=101) at age 14 was associated with reduced ventral striatal activation compared to a sample of 123 healthy subjects matched in age, sex handedness and imaging site (11); furthermore, lower ventral striatal activation at age 14 in healthy adolescents was associated with an increased risk of transition to sub threshold or clinical depression at age 16 in this sample. Whilst anhedonia was associated with reduced ventral striatal activation, accentuated in the presence of both anhedonia and low mood, the presence of low mood without anhedonia was associated with normal ventral striatal reward anticipation. A recent analysis of IMAGEN data showed that 149 individuals with high levels of psychotic like experiences at age 19 had reduced reward related brain activity at age 14 on the MID (12). Taken together, these findings indicate that reduced reward anticipation on the MID task is associated with depression and schizophrenia in adults, and is related to anhedonia, clinical depression status and possibly depression or psychotic-like symptom pathogenesis in mid-adolescence. Whilst the aforementioned studies have shown associations between lower striatal activation during reward anticipation and psychopathology, other studies have suggested that increased activation during reward anticipation may be associated with psychopathology. For example, a recent study (13) showed that greater striatal (caudate) activation during reward anticipation was associated with greater social anxiety in a sample of 40 children age 10-13.

However, it is not yet known at what stage in psychotic or depressive illness do the neural signals of reward anticipation become compromised. No previous large study has used the adult MID task investigate neural reward anticipation signals in early or middle childhood, although “child-friendly” modified versions of the MID task have been shown to engage similar brain regions to the adults MID task (14). Furthermore, no large scale study has examined the relation of reward anticipation to psychopathology in the pre-adolescent stage. A complete understanding of the relationships between reward anticipation and the emergence of psychopathology and mental illness will require examination of precision in the relationships between neutral processing of reward anticipation and the the timeframe of the development of psychopathology in adolescence and young adulthood. A necessary step is to show that the MID engages the classic reward processing network in pre-adolecents and whether or not it relates to psychopathology (especially depression and psychotic-like experiences) before adolescence. Preliminary analysis of the reward receipt phase of the MID task in the large Adolescent Brain Cognitive Development (ABCD) study (15) suggested it produced similar task activations to the those previously seen in adults (16). We therefore used the publicly available ABCD dataset to investigate the following questions:

1. Does the MID task as used in the ABCD study engage similar brain regions during reward anticipation in the pre-adolescent period as previously reported in adolescence and adulthood?
2. Is reduced brain activation in reward anticipation associated with severity or presence of psychotic-like experiences in the pre-adolescent stage?
3. Is reduced brain activation in reward anticipation association with the severity or presence of depressive symptoms in the pre-adolescent stage?
4. Is reduced brain activation in reward anticipation associated with anhedonia in the pre-adolescent stage?

## Methods

### Subjects

3923 children aged 9-11 years from the ABCD dataset (Data Release 1.0) were initially included. These subjects were drawn from 21 centres throughout the US, with participants largely recruited through the school system. Sampling plans and recruitment procedures based on considerations of age, gender, race, socio-economic status and urbanicity were designed to reflect the sociodemographics of the US. Details of recruitment and study design are described elsewhere (17). Details of demographic, physical and mental health assessments are described elsewhere (18). Institutional review board approval was obtained for each site before data collection. All parents provided written informed consent and all children provided assent. Task imaging data was available for 3179 subjects. Subjects with a diagnosis of ADHD, autism spectrum disorder, schizophrenia and intellectual disability, or who were under-weight as defined by CDC, were excluded from analysis, as were subjects who had poor task performance (see below). A total of 2129 subjects were ultimately included in the analysis. Demographic data is described in Table 1.

**Table 1.** Psychopathology in relation to demographics. Psychotic-like experiences were assessed with the Prodromal Questionnaite Child Version, leading to prodromal psychosis scores (PPS). Number of depressive symptoms were measured with K-SADS. Note: Depression symptom scores were unavailable for n = 27 subjects.

|  | PPS |  |  | Depressive Symptoms |  |  |
| --- | --- | --- | --- | --- | --- | --- |
|  | Low (< 7.5) | High (> 7.5) | test for significance | Absent (n = 1956) | Present (n = 146) | test for significance |
| age (months) | 120.3 (7.4) | 118.9 (7.4) | t = 2.5, p = 0.01 | 120 (7.4) | 119.4 (7.8) | t = 1.3, p = 0.2 |
| sex (F / M) | 982 / 963 | 103 / 81 | X<sup>2</sup> = 1.8, p = 0.18 | 988 / 968 | 81 / 65 | X<sup>2</sup> = 1.1, p = 0.23 |
| household income (lower; middle; higher) | 265 / 687 / 870 | 31 / 64 / 67 | X<sup>2</sup> = 3.5, p = 0.17 | 244 / 683 / 899 | 45 / 60 / 27 | X<sup>2</sup> = 60, p < 0.0 |
| parental education (high school; college; post grad) | 207 / 1182 / 556 | 33 / 108 / 43 | X<sup>2</sup> = 9.7, p = 0.01 | 202 / 1185 / 569 | 36 / 90 / 20 | X<sup>2</sup> = 36, p < 0.0 |
| BMI (lean; overweight; obese) | 1374 / 275 / 296 | 136 / 23 / 25 | X<sup>2</sup> = 0.9, p = 0.65 | 1405 / 273 / 278 | 86 / 23 / 37 | X<sup>2</sup> = 15, p < 0.0 |
| Psychopathology Score (mean, sd) | 1.5 (1.9) | 10.5 (2.6) | t = 45.7, p < 0.0 | 0 (0) | 3.1 (2) | t = 18, p < 0.0 |

## Mental Health

### Psychotic-like experiences

Psychotic-like experiences were measured with the prodromal psychosis score (PPS), the sum of the number of yes responses to 21 questions in the Prodromal Questionnaire Child Version (19) (20). These scores were taken as continuous independent predictor variables in the regression model. Across the entire group, mean PPS was 2.3, median = 1, sd = 3.2. The distribution was heavily, positively skewed (see Figure 1). For subjects with PPS > 0, the mean score was 3.9 (sd 3.3, median = 3, range 1 - 21). PPS scores were also binarised at a cut-off of 7.5 for additional regression analysis (n = 181 subjects with higher scores versus remainder of the sample with lower scores).

**Figure 1.**
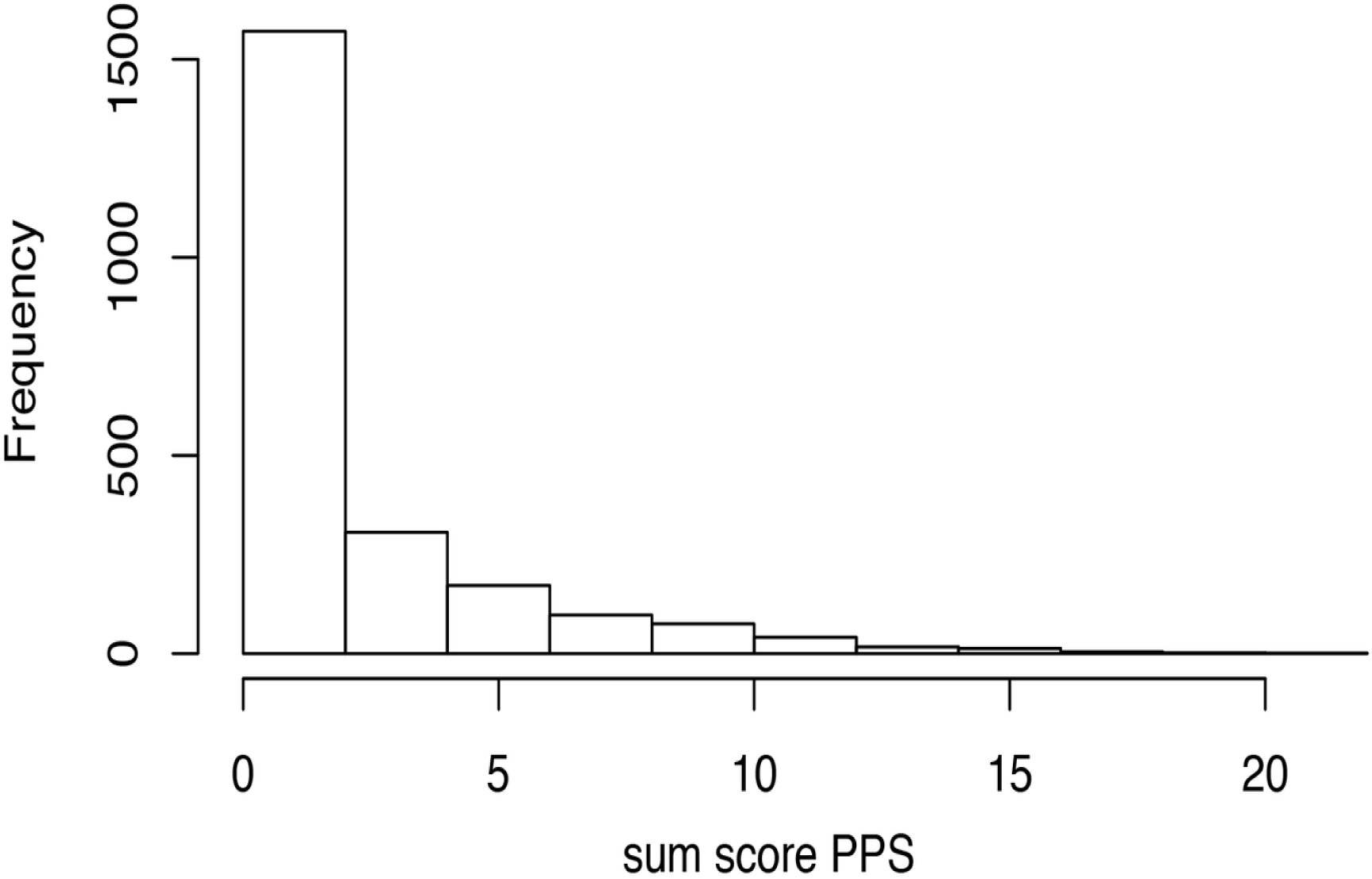
Histogram of sum scores for psychotic like experiences (PPS) for 2129 subjects from the ABCD cohort.

### Depressive symptoms and anhedonia

The number of depressive symptoms present at time of interview were scored based on the youth diagnostic interview K-SADS. Positive symptoms were recorded as present based on affirmative answers to the following: hypersomina, fatigue, concentration disturbance, indecision, decreased appetite, weight loss, increased appetite, weight gain, psychomotor agitation in depressive disorder, psychomotor retardation, guilt, hopelessness, decreased self-esteem, impairment in functioning due to depression, depressed mood, irritability and anhedonia. 146 subjects had at least one or more depressive symptoms. Of these, the mean number of depressive symptoms was 3 (sd 2, median 3, range 1 - 11). Depressive symptom scores were not available for 27 subjects. Depressive symptoms were also binarised (present vs. not present) for additional regression analysis. 75 subjects reported anhedonia.

### Behavioural task in fMRI scanner: Monetary Incentive Delay task

The fMRI Monetary Incentive Delay (MID) task measures domains of reward processing, including anticipation of reward, our interest here. Each trial of the MID task begins with an incentive cue of five possible trial types (Win $.20, Win $5, Lose $0.2, Lose $5, $0 - no money at stake), a delay, a target during which the participant responds to either win money of avoid loosing money and feedback. Each participant receives 40 reward and loss anticipation trials and 20 no money anticipation trials.

The task was programmed in E-Prime professional 2.0 versions 2.0.10.356 or later. The tasks and stimuli are available for download at http::fablab.yale.edu/page/assays-tools. The response collection device was harmonised for precision in response latency across all tasks and all sites with a Current Designs 2-button box. The task was programmed to accept input from the dominant hand. No mandate for precise visual or auditory resolution was imposed across sites.

Task performance was individualised with the initial response target duration based on the participants performance during a practice session prior to scanning. Performance was calculated as the average reaction time (RT) on correct trials plus two standard deviations. To reach a 60% accuracy rate, the task difficulty was adjusted over the course of the task after every third incentivised trial based on the overall accuracy rate of the previous six trials. For feedback, the adaptive algorithm results in 24 positive feedback trials (for both reward and loss) and 16 negative feedback trials (for both reward and loss) on average. Subjects were excluded if participant did not have an acceptable performance in task (this was assessed based on the threshold that all trial types must yield more than 3 events for both positive and negative feedback (n =127)) (indicated in database by mid$beh_mid_perform_flag).

### Imaging Protocols

We used imaging derived phenotype scores provided by the ABCD study team in the data-release 1.0. Data acquisition protocols and analysis protocols for the ABCD dataset are described elsewhere (21) and are reproduced briefly here mainly using wording provided by the ABCD study team. The key events of interest in the current report were the presentation of the cue indicating the potential for a large reward, and the cue indicating no money at stake. We had one contrast of interest in this study, formed by contrasting the brain activation associated with these respective events. Thus the (regional) beta weights for MID anticipation of large reward vs. no money at stake (neutral contrast) were average across runs resulting in the dependent variables. Subjects with MID values above or below 3 standard deviations were removed. Finally 2129 participants were included.

Data acquisition protocol were harmonized for three 3T scanner platforms (Siemens Prisma, General Electric 750 and Philips), used across the 21 data acquisition sites. Scanning sessions occurred across 1 or 2 sessions. The image analysis methods included the following: head motion corrected by registering each frame to the first using AFNI’s 3dvolreg; *B*_0_ distortions were corrected using the reversing gradient methods; displacement field estimated from spin-echo field map scans were applied to gradient - echo images after ajdustment for between-scan head motion; correction for gradient non-linearity distortions; between scan motion correction across all fMRI scans; registration between *T*_2_-weights, spin-echo *B*_0_ calibration scans and *T*_1_-weighted structural images performed using mutual information.

Task-specific pre-processing included removal of initial volumes (Siemens 8TRs; Philips 8TRs; GE DV25 5 TRs; GE DV26 16TRs), normalization and demeaning. Estimation of task-related activation strength used a general linear model (GLM) using AFNIs 3dDeconvolve, with nuisance regressors to model baseline, quadratic trend and motion (motion estimates, derivatives, and squared estimates and derivatives included); time points with frame-wise displacement (FD > 0.9mm) were censored; the hemodynamic response function was modelled as a gamma function with temporal derivatives using AFNIs SPMG model. Events were modeled as instantaneous. The resulting GLM coefficients and t-statistics were sampled on to FreeSurfer-generated cortical surface projected 1mm into cortical grey matter.

The regionally averaged beta weights for the contrast of anticipation of large reward vs. neutral anticipation were taken as the dependent variables (the average across both trial runs). These were further filtered to remove subjects with MID values above or below 3 standard deviations. Finally 2129 participants were included.

### Regional analysis

Processed task data were mapped to 33 cortical regions per hemisphere based on the Deskian-Killany atlas (22) and for subcortical structures for each hemisphere: namely thalamus, caudate, putamen, pallidum, hippocampus, amygdala, accumbens area and ventral diencephalon. Segmentations of these structures were based on FreeSurfer *(aseg) sub-cortical parcellations*. These “imaging derived phenotypes” are provided by the ABCD study team in the data-release. Subjects with activation values above or below 3 standard deviations were excluded. The mean activation per region was subsequently calculated across all subjects. A one-tailed t-test was used to assess difference from 0 and these results were used to create a table and map of regions of significant activation (at *α* < 0.05).

### Statistical Analysis

Multivariate linear regression was used to assess the relationship between PPS, depressive symptoms and fMRI reward anticipation activity. The following additional covariates were included in all analysis; age, sex, race, BMI, household income, parental education, mean frame-wise motion during task. In order to control for variance due to site, we included the variable “device serial number” which indicates the individual MR scanners used to acquire data (n = 25). The number of subjects acquired per scanner varied from a minimum of 9 to a maximum of 283. Site explained a significant (a at 0.05) amount of variance of PPS (F = 7.7, p < 0.00001), but not number of depressive symptoms. False discovery rate (FDR) methods were used to correct results for multiple comparisons across multiple brain regions.

## Results

### Psychotic like experiences - prodromal psychosis scores

PPS were significantly predicted by age (*β* = −0.05, t = −2.3, p = 0.02), but were not predicted by sex, parental education, in-scanner motion or BMI levels. A significant amount of variance in PPS was further predicted by site (F_24,1887_ = 7.4, p < 0.001), household income (F_2,1887_ = 7.9, p < 0.001) and race/ethnicity (F_3,1887_ = 5.4, p = 0.001).

### Number of depressive symptoms

Number of depressive symptoms was significantly predicted by household income (lower vs. middle *β* = −0.18, t = −2.4, p = 0.02; lower vs. higher *β* = −0.3, t = −3.7, p =0.0003) and BMI levels (lean vs. overweight *β* = −0.01, t = −0.17, p = 0.87; lean vs. obese *β* = 0.19, t = 2.9, p = 0.004).

### Mean reward anticipation activation

Results of mean activation for top 10 brain regions are reported in Table 2 (full list available in Supplementary Material).

**Table 2.** Mean activation and results of one-tailed t-test for significance for top 10 cortical regions / sub-cortical structures. All p-values less than 0.001

| Cortical region / sub-cortical structure | mean activation | t-stat | p-value | FDR-corrected p-value |
| --- | --- | --- | --- | --- |
| right caudate | 0.11 | 21.35 | < 0.0 | < 0.0 |
| left caudate | 0.1 | 19.12 | < 0.0 | < 0.0 |
| left accumbens area | 0.1 | 18.54 | < 0.0 | < 0.0 |
| right putamen | 0.07 | 16.9 | < 0.0 | < 0.0 |
| left putamen | 0.07 | 16.07 | < 0.0 | < 0.0 |
| right accumbens area | 0.09 | 15.67 | < 0.0 | < 0.0 |
| left pallidum | 0.06 | 14.42 | < 0.0 | < 0.0 |
| right thalamus proper | 0.05 | 14.15 | < 0.0 | < 0.0 |
| right caudal anterior cingulate | 0.06 | 14.06 | < 0.0 | < 0.0 |
| right ventral diencephalon | 0.06 | 14.03 | < 0.0 | < 0.0 |

A plot of mean activation per cortical region, averaged across 2129 subjects is reported below (Figure 2). Results are filtered to exclude regions where activation did not significantly differ from 0 (at *α* = 0.05).

**Figure 2.**
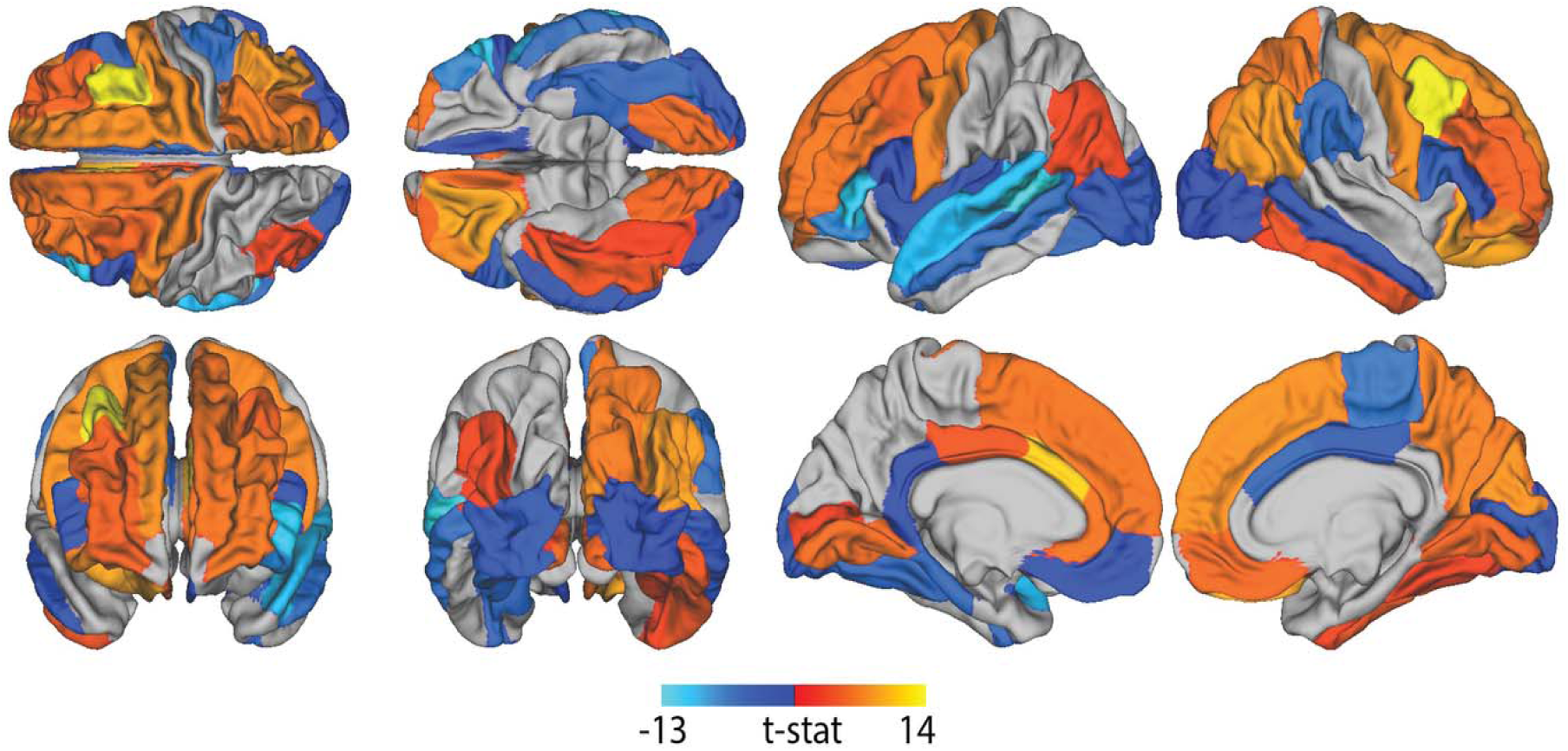
Cortical regions of statistically significant reward anticipation activation (as assessed from one-tailed t-test in 2129 subjects).

### Regression of PPS on MID task activation

After FDR correction, PPS as a continuous variable was not found to be a significant predictor of MID in any brain region. PPS was predictive of MID activation in the right parsorbitalis region (*β* = −0.07, t = −2.8, p = 0.01) and for the left accumbens area (*β* = −0.06, t = −2.34, p = 0.02), however these did not survive correction for multiple comparisons (Table 3).

**Table 3.** Results of regression of PPS on MID task activation (top 10 regional associations).

| Cortical region / sub-cortical structure | PPS (continuous) |  |  |  | PPS (binarised) |  |  |  |
| --- | --- | --- | --- | --- | --- | --- | --- | --- |
|  | beta | t-stat | p-value | FDR-corrected p-value | beta | t-stat | p-value | FDR-corrected p-value |
| right parsorbitalis | -0.07 | -2.8 | 0.01 | 0.42 | -0.3 | -3.53 | 0.0 | 0.04 |
| left accumbens area | -0.06 | -2.34 | 0.02 | 0.8 | -0.15 | -1.79 | 0.07 | 0.94 |
| left inferior parietal | -0.04 | -1.58 | 0.11 | 0.97 | -0.14 | -1.69 | 0.09 | 0.94 |
| right hippocampus | 0.03 | 1.44 | 0.15 | 0.97 | 0.04 | 0.42 | 0.67 | 0.97 |
| right caudate | -0.03 | -1.42 | 0.16 | 0.97 | -0.11 | -1.27 | 0.21 | 0.97 |
| right lateral orbitofrontal | -0.03 | -1.38 | 0.17 | 0.97 | -0.16 | -1.9 | 0.06 | 0.94 |
| left pericalcarine | -0.03 | -1.36 | 0.17 | 0.97 | -0.08 | -0.89 | 0.37 | 0.97 |
| left lateral orbitofrontal | -0.03 | -1.28 | 0.2 | 0.97 | -0.19 | -2.28 | 0.02 | 0.92 |
| left cuneus | -0.03 | -1.15 | 0.25 | 0.97 | -0.11 | -1.28 | 0.2 | 0.97 |
| left caudal middle frontal | -0.03 | -1.1 | 0.27 | 0.97 | -0.13 | -1.58 | 0.11 | 0.94 |

When PPS was binarised, there was a significant difference between subjects with higher scores compared to lower in terms of predicting MID task activation in the right parsorbitalis (higher vs. lower *β* = −0.3, t = −3.5, p = 0.04) after FDR correction, but no other significant associations. A list of full results is available in Supplementary Material.

### Regression of number of depression symptoms on MID task activation

Without correction for multiple comparisons, number of depression symptoms was found to be predictive of MID reward anticipation task activity in the left ventral diencephalon and several cortical regions, namely the fusiform in the right hemisphere and the temporal pole, fusiform, middle temporal and superior parietal cortex in the left hemisphere (Table 4). However none of these results survived correction for multiple comparisons.

**Table 4.** Results of regression of (a) number of depressive symptoms and (b) presence of depressive on reward anticipation activation (top regional associations).

| Cortical region / sub-cortical structure | No. depression symptoms |  |  |  | Depressive symptoms present |  |  |  |
| --- | --- | --- | --- | --- | --- | --- | --- | --- |
|  | beta | t-stat | p-value | FDR-corrected p-values | beta | t-stat | p-value | FDR-corrected p-values |
| right fusiform | 0.06 | 2.78 | 0.01 | 0.23 | 0.19 | 2.05 | 0.04 | 0.98 |
| left temporal pole | 0.06 | 2.77 | 0.01 | 0.23 | 0.18 | 1.88 | 0.06 | 0.98 |
| left ventral dc | 0.06 | 2.46 | 0.01 | 0.32 | 0.18 | 1.89 | 0.06 | 0.98 |
| left fusiform | 0.06 | 2.42 | 0.02 | 0.32 | 0.13 | 1.35 | 0.18 | 0.98 |
| left middle temporal | 0.05 | 2.12 | 0.03 | 0.57 | 0.05 | 0.52 | 0.6 | 0.98 |
| left superior parietal | 0.05 | 1.98 | 0.05 | 0.66 | 0.08 | 0.81 | 0.42 | 0.98 |
| right middle temporal | 0.04 | 1.84 | 0.07 | 0.78 | 0.06 | 0.6 | 0.55 | 0.98 |
| left amygdala | 0.04 | 1.64 | 0.1 | 0.84 | 0.04 | 0.41 | 0.68 | 0.98 |
| left transverse temporal | -0.04 | -1.63 | 0.1 | 0.84 | -0.2 | -2.13 | 0.03 | 0.98 |
| right superior parietal | 0.04 | 1.62 | 0.11 | 0.84 | 0.04 | 0.42 | 0.68 | 0.98 |

When number of depression symptoms were binarised for those with vs. without symptoms and regression repeated, no regions were significantly predictive of MID task activation after FDR-correction (Table 4).

When this analysis was repeated for the presence of anhedonia (n = 75), we found that the presence of anhedonia significantly (positively) predicted MID task activation in the left temporal pole, the right fusiform and (negatively) the right parsorbitalis (Table 5); however these results did not survive correction for multiple comparisons.

**Table 5.**
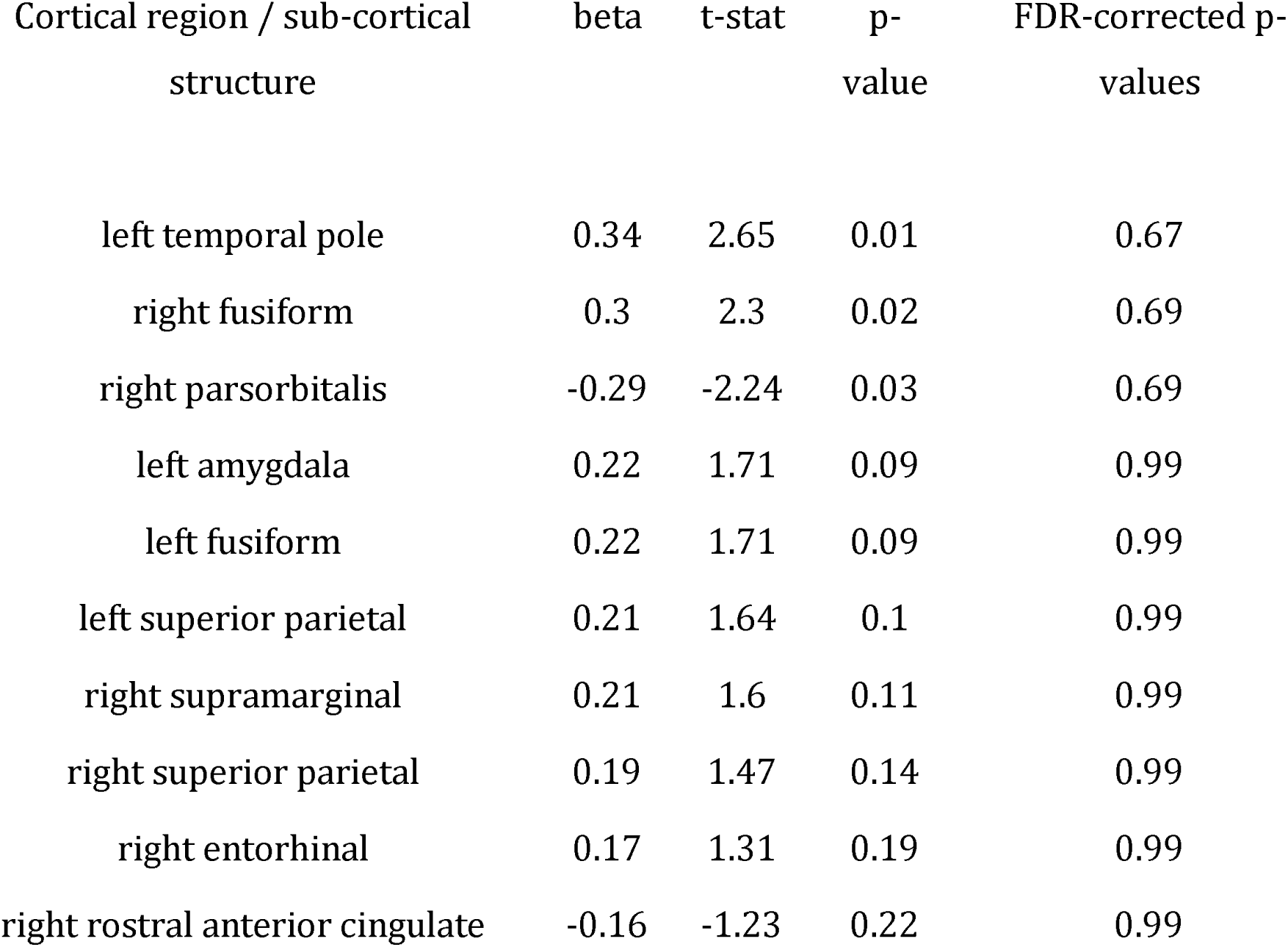
Results of regression of presence of anhedonia on reward anticipation activation (top 10 regional associations).

## Discussion

In this large study of 9-10 year olds, the neural correlates reward anticipation were similar to those previously observed in older adolescents and adults, extending the preliminary results of reward feedback in this cohort (16). There was, however, little evidence that individual differences in brain reward anticipation signals were associated with severity of psychotic-like experiences or number of depressive symptoms. There were no linear associations between brain reward anticipation and severity of psychotic-like experiences or number of depressive symptoms that survived correction for multiple comparison. There were a handful of associations that had p-values below 0.05 without multiple comparison correction, but none survived correction for multiple comparison. For example there were (uncorrected) negative associations between activations in left accumbens and right orbitofrontal cortex with psychotic like experiences; these associations was only present in the left, not right, accumbens, and the right not left orbitoftonal cortex, and are likely due to chance. There was a negative association between right orbitofrontal cortex activation and presence of anhedonia, and positive association between temporal lobe activations and presence of anhedonia, and between temporal lobe (temporal pole, fusiform gyrus, left middle temporal gyrus) activations number of depressive symptoms. None of these associations was significant after correction for multiple comparisons. Our results indicate that variations in brain reward anticipation signals at age 9-10 are not associated with the severity of psychotic-like experiences, or number of depressive symptoms.

There was a solitary finding that survived corrected for multiple comparison for the number of brain regions studied: namely children with presence of psychotic-like experiences had less activation in the right parsorbitalis (orbital aspect of inferior frontal gyrus) compared to those without such experiences. This isolated finding only in one hemisphere must be considered preliminary, as although we corrected for the number of brain regions studied, we repeated the analyses with several different psychiatric phenotypes and did not correct for these multiple phenotypic analyses. Furthermore, although a plausible post-hoc rationale could be constructed for the importance of the right parasobitalis in the genesis of psychotic like experiences, it does not correspond to the bulk of the prior findings in the literature relating reward anticipation to psychotic illness or psychotic psychopathology in older groups.

The task used in the MID study engaged similar brain regions during reward anticipation as have previously been seen in adults and adolescents in very similar versions of the same task. Thus, we can eliminate the possibility that this largely null result with psychopathology was linked to fMRI measurement failure or developmental factors related to the developmental trajectory of the function of regions/structures representing reward anticipation. The possibility of measurement error in terms of psychopathology at age 9-10 could be considered, but is unlikely to be solely responsible in our view. Psychotic-like experiences were measured with a new scale developed especially for this study, which has been shown to perform well in this sample (19). Depression was measured with the K-SADS, the gold standard in this age group for diagnostic purposes; admittedly, in terms of a continuous measures of severity of depression, or of anhedonia, additional assessments might be more sensitive to individual differences in participants. Given that in middle adolescence, and in adulthood, striatal reward anticipation has been shown to be deficient in patients with schizophrenia and clinical or subthreshold depression (4)(5)(6)(11)(12), it appears that clear associations between psychopathology and brain reward anticipation signatures only emerge during adolescence. The ABCD study provides a platform through which this prediction can be tested, as the study protocol is that repeat measures of fMRI and psychopathology will be acquired in the coming years.

The significance of psychotic-like experiences in the general population is nuanced (23). Psychotic-like experiences at the age of around 10 years are relatively common (around 17%) (24; 25). Whilst their presence indicates increased risk of current or subsequent mental disorder, only a minority of children with psychotic-like experiences at this age will go on to develop a psychotic illness (23;24). Thus the presence of such symptoms should not be seen as, in isolation, marking the presence of mental illness or of specific schizophrenia spectrum risk However, the presence of such symptoms does indicate a raised risk (compared to children who do not express psychotic like experiences) of future mental disorder, including schizophrenia spectrum disorder, of future anxiety and depressive disorders, and of suicidal thoughts and behaviours. The frequency of endorsement of psychotic like expeirences declines over the adolescent period, but the strength of the association with mental disorder strengthens in this period for those in whom psychotic experiences persist. Thus, as age 10-11, an endorsement of psychotic like experiences items on interview is less likely to be representative of mental disorder than a similar endorsement in adolescence (26), and the persistence of psychotic-like experiences over time has been shown to be associated with a higher risk of mental disorder than isolated measurements (24; 25). Previous studies have suggested links between reward processing (and other fMRI task) activation and psychopathology in children and adolescents than the current sample (11)(27; 12). This leads to the following prediction: children who persist in endorsing psychotic-like experiences at multiple time-points over the coming years will be more prone to develop schizophrenia spectrum or depressive illnesses than other short members, and this persistence of psychopathology, and emergence of new psychopathology in follow-ups, may be associated with (or possibly predicted by) concomitant deficits in brain activation in the MID task. This prediction can be tested in follow-up waves of the ABCD study.

The study has a number of limitations. We used various exclusion criteria including diagnosis of ADHD, which affects several hundred children in the cohort. ADHD has been associated with abnormal brain reward processing previously as well as various psychiatric symptoms such as depressive symptoms and psychotic symptoms; furthermore ADHD is commonly treated with stimulant medication that may alter brain activation during reward anticipation, making results challenging to interpret. Thus, excluding ADHD is helpful in reducing the chances of finding spurious associations between various domains of psychopathology and altered brain reward anticipation signals; however, it reduces the sample size, thus reducing statistical power, and it makes the sample less representative of the general population. Additional children had to be excluded for other reasons, such as artefacts, movement in the scanner or non-engagement with the task. The study data were collected on multiple MRI scanners, which is likely an additional source of noise and variance. We try to compensate for this by taking scanner site into account in statistical analyses. Acquiring data at a single site using a single scanner could be preferable, in terms of reducing sources of variability; however, a major advantage of a multi-site study is the potential for much larger sample size.

There are many measures of potential interest that can be generated from the MID task. We calculated reward anticipation by contrasting anticipation of a large reward versus no reward was it is an effective and reliable contrast in eliciting ventral striatal and medial prefrontal reward anticipation associated activation. We used this same contrast in our previous MID study (6) as did Jia et al (2015) in IMAGEN, the largest MID study previously (10). Many other related contrasts could have been used simply to probe reward anticipation, and the task data could also or alternatively be used to examine anticipation of punishment, brain responses to reward or punishment receipt or omission of expected rewards and/or punishments; numerous connectivity analyses are also possible. Whilst these alternative analyses would be of considerable interest, each additional analysis increases the liklihood of type I error unless rigorous correction for multiple comparison is made, in which case the likelihood of type II error is increased. It is possible that a voxel-wise or vertex-wise analysis could be more sensitive than the approach we used, of regionally averaged scores. However, we note that the regionally averaged scores revealed highly significant task activation in the expected areas of striatal subdivisions, thalamus and medial frontal cortex, supporting their validity.

In summary, in spite of the MID task eliciting robust subcortical and cortical activation in a large sample of 9-10 year olds, there were no strong associations with severity of psychopathology (psychotic-like experiences or depressive or anxiety symptoms) in this age group. It will be of considerable interest to examine whether baseline activation in this age is predictive of the development of future psychopathology, or whether abnormal patterns of brain activation emerge during the maturation process, perhaps associated with concomitant development of specific of general measures of psychopathology.

## Author roles

LR.: Conceptualization, methodology, formal analysis, writing (original draft preparation, review and editing); GKM. Conceptualization, methodology, writing (original draft preparation, review and editing).

## Acknowledgments

We thank the ABCD study team and participants, and Nicole Karcher for advice on K-SADS scoring.

## Supplementary Material

### Origin of raw data

Diagnoses from data table “abcd_screen01.csv”

Age, sex and BMI from “abcd_ANT01”

Household income, race and parental education from “abcd_pdem01”

Scanner (device) number from “abcd_mristv01”

Depression, PPS diagnosis from “abcd_ksad501” (PPS: pps_y_ss_number)

Motion from “abcd_midaparc01” (fmri_beta_gparc_mean_motion)

Performance in scanner from “abcd_mid01” (beh_mid_perform_flag).

MID data from “abcd_midaparc01” (ASEG = “mdaclgrwvsntb”; APARC=).

Prodromal psychosis scale “abcd_pps01”.

### Supplementary Results: Mean Activation

*Mean activation for all cortical regions (n = 66) and sub-cortical structures (n = 16), including results of one-tailed t-tests for significance*.

| Cortical region / sub-cortical structure | mean activation | t-stat | p-value |
| --- | --- | --- | --- |
| right caudate | 0.11 | 21.35 | 1.03e-91 |
| left accumbens area | 0.1 | 18.54 | 3.69e-71 |
| left caudate | 0.1 | 19.12 | 2.81e-75 |
| right accumbens area | 0.09 | 15.67 | 1.98e-52 |
| right putamen | 0.07 | 16.9 | 3.4e-60 |
| left putamen | 0.07 | 16.07 | 6.6e-55 |
| left ventral dc | 0.07 | 13.84 | 9.22e-42 |
| right caudal anterior cingulate | 0.06 | 14.06 | 5.74e-43 |
| right ventral dc | 0.06 | 14.03 | 8.54e-43 |
| left pallidum | 0.06 | 14.42 | 5.11e-45 |
| right pallidum | 0.06 | 13.79 | 1.76e-41 |
| right thalamus proper | 0.05 | 14.15 | 1.69e-43 |
| right lateral occipital | 0.05 | 8.54 | 2.45e-17 |
| left caudal anterior cingulate | 0.05 | 10.88 | 7.44e-27 |
| left thalamus proper | 0.04 | 11.31 | 8.19e-29 |
| right rostral middle frontal | 0.04 | 7.02 | 3.08e-12 |
| right fusiform | 0.04 | 7.82 | 8.18e-15 |
| left lingual | 0.03 | 4.85 | 1.3e-06 |
| right posterior cingulate | 0.03 | 7.13 | 1.4e-12 |
| right lingual | 0.03 | 4.64 | 3.67e-06 |
| right lateral orbitofrontal | 0.03 | 4.45 | 8.84e-06 |
| left rostral middle | 0.03 | 5.47 | 4.92e-08 |
| right parsopercularis | 0.03 | 6.8 | 1.32e-11 |
| right caudal middle frontal | 0.03 | 7.23 | 6.62e-13 |
| left rostral anterior cingulate | 0.03 | 5.3 | 1.27e-07 |
| right entorhinal | 0.03 | 2.22 | 0.03 |
| left precentral | 0.02 | 6.49 | 1.06e-10 |
| right precentral | 0.02 | 6.12 | 1.1e-09 |
| right rostral anterior cingulate | 0.02 | 4.22 | 2.53e-05 |
| right superior frontal | 0.02 | 5.77 | 9.14e-09 |
| left superior frontal | 0.02 | 5.31 | 1.19e-07 |
| left pericalcarine | 0.02 | 2.22 | 0.03 |
| left caudal middle | 0.02 | 4.29 | 1.88e-05 |
| right pericalcarine | 0.02 | 2.0 | 0.05 |
| left posterior cingulate | 0.01 | 3.41 | 0.0 |
| right inferior parietal | 0.01 | 3.42 | 0.0 |
| left inferior parietal | 0.01 | 2.51 | 0.01 |
| left precuneus | -0.01 | -2.07 | 0.04 |
| left hippocampus | -0.01 | -2.29 | 0.02 |
| left pars opercularis | -0.01 | -2.28 | 0.02 |
| left isthmuscingulate | -0.01 | -2.36 | 0.02 |
| right isthmuscingulate | -0.01 | -2.28 | 0.02 |
| right postcentral | -0.02 | -3.58 | 0.0 |
| right paracentral | -0.02 | -4.24 | 2.35e-05 |
| right parstriangularis | -0.02 | -3.48 | 0.0 |
| right bankssts | -0.02 | -4.63 | 3.84e-06 |
| right superior temporal | -0.02 | -5.29 | 1.37e-07 |
| left lateral occipital | -0.02 | -3.38 | 0.0 |
| left meidal orbitofrontal | -0.02 | -2.41 | 0.02 |
| left fusiform | -0.02 | -4.85 | 1.33e-06 |
| left parahippocampal | -0.02 | -4.9 | 1.01e-06 |
| right amygdala | -0.02 | -4.35 | 1.43e-05 |
| left middle temporal | -0.03 | -5.41 | 6.87e-08 |
| right parahippocampal | -0.03 | -5.76 | 9.39e-09 |
| left temporal pole | -0.03 | -2.76 | 0.01 |
| left amygdala | -0.03 | -6.06 | 1.62e-09 |
| right parsorbitalis | -0.03 | -2.93 | 0.0 |
| right medial orbitofrontal | -0.03 | -2.99 | 0.0 |
| left superior temporal | -0.03 | -9.49 | 6.13e-21 |
| left frontal pole | -0.04 | -2.17 | 0.03 |
| left bankssts | -0.05 | -12.55 | 6.75e-35 |
| left parstriangularis | -0.06 | -10.75 | 2.75e-26 |
| left parsorbitalis | -0.07 | -6.8 | 1.4e-11 |

### Supplementary Results: psychotic-like symptoms and reward anticipation activation

Results of regression of PPS on reward anticipation activation for all cortical regions / sub-cortical structures. Note p-values are not FDR-corrected.

| Region ID | Cortical region / sub-cortical structure | beta | t-stat | p-value |
| --- | --- | --- | --- | --- |
| 67 | right parsorbitalis | -0.07 | -2.8 | 0.01 |
| 7 | left accumbens area | -0.06 | -2.34 | 0.02 |
| 23 | left inferior parietal | -0.04 | -1.58 | 0.11 |
| 13 | right hippocampus | 0.03 | 1.44 | 0.15 |
| 10 | right caudate | -0.03 | -1.42 | 0.16 |
| 60 | right lateral orbitofrontal | -0.03 | -1.38 | 0.17 |
| 36 | left pericalcarine | -0.03 | -1.36 | 0.17 |
| 27 | left lateral orbitofrontal | -0.03 | -1.28 | 0.2 |
| 20 | left cuneus | -0.03 | -1.15 | 0.25 |
| 19 | left caudal middle frontal | -0.03 | -1.1 | 0.27 |
| 75 | right rostral middle temporal | -0.02 | -1.0 | 0.32 |
| 78 | right superior temporal | 0.02 | 0.99 | 0.32 |
| 63 | right middle temporal | 0.02 | 0.96 | 0.34 |
| 5 | left hippocampus | 0.02 | 0.95 | 0.34 |
| 6 | left amygdala | 0.02 | 0.94 | 0.35 |
| 42 | left rostral middle frontal | -0.02 | -0.93 | 0.35 |
| 2 | left caudate | -0.02 | -0.93 | 0.35 |
| 72 | right precentral | -0.02 | -0.92 | 0.36 |
| 69 | right pericalcarine | -0.02 | -0.88 | 0.38 |
| 49 | left transverse temporal | 0.02 | 0.88 | 0.38 |
| 29 | left middle orbitofrontal | -0.02 | -0.86 | 0.39 |
| 21 | left entorhinal | 0.02 | 0.86 | 0.39 |
| 53 | right cuneus | -0.02 | -0.85 | 0.4 |
| 38 | left posterior cingulate | 0.02 | 0.84 | 0.4 |
| 22 | left fusiform | 0.02 | 0.78 | 0.43 |
| 52 | right caudal middle frontal | -0.02 | -0.78 | 0.44 |
| 46 | left supramarginal | -0.02 | -0.76 | 0.45 |
| 8 | left ventral dc | 0.02 | 0.75 | 0.45 |
| 65 | right paracentral | -0.02 | -0.75 | 0.46 |
| 32 | left paracentral | -0.02 | -0.7 | 0.48 |
| 45 | left superior temporal | 0.02 | 0.66 | 0.51 |
| 48 | left temporal pole | 0.02 | 0.63 | 0.53 |
| 82 | right transverse temporal | 0.02 | 0.61 | 0.54 |
| 25 | left isthmuscingulate | 0.01 | 0.6 | 0.55 |
| 68 | right parstriangularis | -0.01 | -0.6 | 0.55 |
| 54 | right entorhinal | 0.01 | 0.56 | 0.57 |
| 43 | left superior frontal | -0.01 | -0.55 | 0.58 |
| 37 | left postcentral | 0.01 | 0.54 | 0.59 |
| 64 | right parahippocampal | 0.01 | 0.54 | 0.59 |
| 76 | right superior frontal | -0.01 | -0.53 | 0.59 |
| 55 | right fusiform | -0.01 | -0.49 | 0.62 |
| 11 | right putamen | -0.01 | -0.48 | 0.63 |
| 28 | left lingual | -0.01 | -0.46 | 0.65 |
| 44 | left superior parietal | 0.01 | 0.45 | 0.65 |
| 15 | right accumbens area | -0.01 | -0.45 | 0.65 |
| 81 | right temporal pole | -0.01 | -0.45 | 0.66 |
| 12 | right pallidum | -0.01 | -0.44 | 0.66 |
| 66 | right parsopercularis | -0.01 | -0.43 | 0.67 |
| 40 | left precuneus | -0.01 | -0.43 | 0.67 |
| 80 | right frontal pole | 0.01 | 0.43 | 0.67 |
| 26 | left lateral occipital | 0.01 | 0.42 | 0.67 |
| 3 | left putamen | -0.01 | -0.37 | 0.71 |
| 51 | right caudal anterior cingulate | 0.01 | 0.34 | 0.73 |
| 47 | left frontal pole | -0.01 | -0.34 | 0.74 |
| 74 | right rostral anterior cingulate | 0.01 | 0.34 | 0.74 |
| 70 | right postcentral | -0.01 | -0.33 | 0.74 |
| 41 | left rostral anterior cingulate | 0.01 | 0.33 | 0.75 |
| 33 | left parsopercularis | -0.01 | -0.3 | 0.77 |
| 77 | right superior parietal | -0.01 | -0.3 | 0.77 |
| 39 | left precentral | -0.01 | -0.29 | 0.77 |
| 24 | left inferior temporal | 0.01 | 0.29 | 0.77 |
| 4 | left pallidum | -0.01 | -0.28 | 0.78 |
| 34 | left parsorbitalis | 0.01 | 0.26 | 0.8 |
| 18 | left caudal anterior cingulate | 0.01 | 0.22 | 0.82 |
| 50 | right bankssts | 0.0 | 0.2 | 0.84 |
| 71 | right posterior cingulate | 0.0 | 0.19 | 0.85 |
| 31 | left parahippocampal | 0.0 | 0.19 | 0.85 |
| 30 | left middle temporal | 0.0 | 0.16 | 0.87 |
| 58 | right isthmuscingulate | 0.0 | 0.16 | 0.88 |
| 14 | right amygdala | 0.0 | 0.15 | 0.88 |
| 9 | right thalamus proper | -0.0 | -0.14 | 0.89 |
| 62 | right medial orbitofrontal | -0.0 | -0.13 | 0.89 |
| 17 | left bankssts | 0.0 | 0.13 | 0.9 |
| 1 | left thalamus proper | 0.0 | 0.11 | 0.91 |
| 73 | right precuneus | -0.0 | -0.1 | 0.92 |
| 16 | right ventral dc | 0.0 | 0.1 | 0.92 |
| 61 | right lingual | -0.0 | -0.09 | 0.93 |
| 57 | right inferior temporal | 0.0 | 0.08 | 0.94 |
| 56 | right inferior parietal | 0.0 | 0.06 | 0.95 |
| 79 | right supramarginal | -0.0 | -0.05 | 0.96 |
| 35 | left parstriangularis | 0.0 | 0.03 | 0.97 |
| 59 | right lateral occipital | 0.0 | 0.03 | 0.97 |

